# Accessible, realistic genome simulation with selection using stdpopsim

**DOI:** 10.1101/2025.03.23.644823

**Authors:** Graham Gower, Nathaniel S. Pope, Murillo F. Rodrigues, Silas Tittes, Linh N. Tran, Ornob Alam, Maria Izabel A. Cavassim, Peter D. Fields, Benjamin C. Haller, Xin Huang, Ben Jeffrey, Kevin Korfmann, Christopher C. Kyriazis, Jiseon Min, Ińes Rebollo, Clara T. Rehmann, Scott T. Small, Chris C. R. Smith, Georgia Tsambos, Yan Wong, Yu Zhang, Christian D. Huber, Gregor Gorjanc, Aaron P. Ragsdale, Ilan Gronau, Ryan N. Gutenkunst, Jerome Kelleher, Kirk E. Lohmueller, Daniel R. Schrider, Peter L. Ralph, Andrew D. Kern

## Abstract

Selection is a fundamental evolutionary force that shapes patterns of genetic variation across species. However, simulations incorporating realistic selection along heterogeneous genomes in complex demographic histories are challenging, limiting our ability to benchmark statistical methods aimed at detecting selection and to explore theoretical predictions. stdpopsim is a community-maintained simulation library that already provides an extensive catalog of species-specific population genetic models. Here we present a major extension to the stdpopsim framework that enables simulation of various modes of selection, including background selection, selective sweeps, and arbitrary distributions of fitness effects (DFE) acting on annotated subsets of the genome (for instance, exons). This extension maintains stdpopsim’s core principles of reproducibility and accessibility while adding support for species-specific genomic annotations and published DFE estimates. We demonstrate the utility of this framework by comparing methods for demographic inference, DFE estimation, and selective sweep detection across several species and scenarios. Our results demonstrate the robustness of demographic inference methods to selection on linked sites, reveal the sensitivity of DFE-inference methods to model assumptions, and show how genomic features, like recombination rate and functional sequence density, influence power to detect selective sweeps. This extension to stdpopsim provides a powerful new resource for the population genetics community to explore the interplay between selection and other evolutionary forces in a reproducible, user-friendly framework.

## Introduction

Selection is a fundamental force in evolution, shaping the genetic diversity of populations and driving the adaptation of species to their environments. The effects of selection on genetic variation are complex, and can be difficult to disentangle from other evolutionary processes such as mutation, recombination, and genetic drift (e.g., Ohta, 1973; Lewontin and Krakauer, 1973; Kimura, 1985; Tajima, 1989; Gillespie, 1991; Kern and Hahn, 2018). For instance, changes in population size can lead to fluctuations in genetic diversity across a recombining chromosome that can mimic the effects of selection (Simonsen, Churchill, and Aquadro, 1995), leading to spurious inferences about the strength and targets of genetic adaptation (Simonsen, Churchill, and Aquadro, 1995; Akey et al., 2004; Nielsen et al., 2005). In turn, selection can confound our ability to infer demographic history from allele frequencies (Ewing and Jensen, 2016; Schrider, Shanku, and Kern, 2016) and estimates of inverse coalescent rate (Schrider, Shanku, and Kern, 2016; Johri, Riall, et al., 2021; Cousins et al., 2024). Thus, it is imperative to jointly account for the effects of selection and demography when inferring evolutionary history from genetic data (e.g., Sheehan and Song, 2016; Johri, Charlesworth, and Jensen, 2020), yet the number of tools for doing so remain limited.

To meet the growing need for the interpretation, analysis, and exploration of realistic and complex evolutionary models, the field of population genetics has increasingly turned to simulation. Simulation in population genetics has a long history including both backward-in-time coalescent simulations (Kingman, 1982; Hudson, 1983; Hudson et al., 1990) and forward-in-time simulations of complex demography and selection (e.g., Gillespie, 1984; Hernandez, 2008; Thornton, 2014; Haller and Messer, 2019; Gaynor, Gorjanc, and Hickey, 2021). The development of simulation tools has been driven by several needs: understanding the effects of complex evolutionary processes on genetic variation (e.g., Peischl et al., 2013; Thornton, 2019; Galloway, Cresko, and Ralph, 2020; Zhang, Kim, Lohmueller, et al., 2020), obtaining a null model for hypothesis testing (e.g., Hudson, Boos, and Kaplan, 1992; Hudson, Bailey, et al., 1994; Sabeti et al., 2002), exploring the power and limitations of statistical methods (e.g., Przeworski, 2002; MacLeod, Hayes, and Goddard, 2014), and increasingly, providing a basis for machine learning and other simulation-based inference methods (e.g., Beaumont, Zhang, and Balding, 2002; Lin et al., 2011; Kern and Schrider, 2018; Mughal and DeGiorgio, 2019; Sanchez et al., 2021; Wang et al., 2021; Zhang, Kim, Singh, et al., 2023; Smith and Kern, 2023). However, joint simulation of complex demography and selection on realistic genomes is challenging, requiring many choices including the strength and rate of selection, and a parameterized demographic model, as well as implementation of genetic maps and other genomic features related to the strength of selection. This is a daunting task, and can be a barrier to the adoption of simulation-based methods in population genetics. Furthermore, different simulation and inference tools often use different structures and parameter scalings for their models, making it difficult to directly compare across these tools and the studies that use them.

In light of the complexities associated with estimating and simulating population genetic models, it is no surprise that a lingering challenge with simulation in population genetics has been reproducibility and the ability to share and compare results among researchers (e.g., Ragsdale, Nelson, et al., 2020). This challenge has been addressed in part by the development of community resources for sharing and distributing simulation software via the stdpopsim project (Adrion et al., 2020). stdpopsim provides a standardized interface for accessing a wide range of population genetic models, and has been widely adopted by the community (e.g., Speidel et al., 2021; Wang et al., 2021; Yang et al., 2022; DeGiorgio and Szpiech, 2022; Browning, Waples, and Browning, 2023; Schweiger and Durbin, 2023; Temple, Waples, and Browning, 2024; Haag, Jordan, and Stamatakis, 2025). Yet, previous releases of stdpopsim did not include models of selection, which is a major limitation for more empirically motivated applications of population genetic simulation. In particular, modeling selection through simulation is critical for understanding processes such as adaptation (e.g. Thornton, 2019; Hartfield and Gĺemin, 2024), the effects of selective sweeps (e.g. Braverman et al., 1995; Fay and Wu, 2000; Przeworski, 2002; Przeworski, Coop, and Wall, 2005; Schrider, Mendes, et al., 2015), and the impact of background selection on genetic diversity (e.g. Charlesworth, Morgan, and Charlesworth, 1993; Charlesworth, Charlesworth, and Morgan, 1995; Williamson and Orive, 2002; Ewing and Jensen, 2016; Torres et al., 2020). Ideally, one would like to have a simple, unified framework for simulating both neutral and non-neutral evolutionary processes, and to be able to compare the results of these simulations to empirical data in a manner that is both accessible to a wide range of researchers and highly reproducible. Furthermore, the framework should include the complex realities of genomes, including heterogeneous recombination rates, variation in the size and density of functional elements, and non-equilibrium demographic histories. Moreover, one should be able to model all of these features simultaneously using estimates and annotations obtained from a species of interest (e.g. Schrider, 2020; Rodrigues, Kern, and Ralph, 2024). stdpopsim provides a framework for such models, thus lowering the barrier to production of complex, multifaceted simulations, and allowing a wider range of researchers to make use of high-quality simulations in developing or testing their work. A central goal of the stdpopsim project is open, collaborative, and inclusive work; we view capacity-building across the community as equal in importance to software development.

Here, we provide an overview of a major new addition to stdpopsim—the inclusion of models of selection. We begin by describing the models of selection that we have implemented, as well as the parameter choices that are available to the user. This includes models of background selection, selective sweeps, and models of the distribution of fitness effects (DFE). Further, we describe how these can be applied not only genomewide but to genomic regions with a particular annotation, specific to a given species available through the stdpopsim application programming interface (API), and combined with a parameterized model of demography and recombination to provide a realistic simulation of genetic variation in a population. We then provide a series of examples of how these models can be used to benchmark and compare the performance of different methods for using genetic data to infer demographic history, the distribution of fitness effects, and the targets of recent positive selection. The examples use genomes of both humans (*Homo sapiens*) and the critically endangered Vaquita porpoise (*Phocoena sinus*), which provides an interesting comparison due to its small recent population size.

### Implementing Selection in stdpopsim

stdpopsim previously provided a wide range of models of neutral demographic history from roughly two dozen species (Lauterbur et al., 2023). This was accomplished through the use of a standardized interface and data structure that allows for the curated addition of new species and models, and can use either msprime (Baumdicker et al., 2021) or SLiM (Haller and Messer, 2019) as the backend engine for efficient population genetic simulation. To extend stdpopsim’s functionality to include models of selection, we introduce two new classes to the stdpopsim API: the DFE class and the Annotation class. The DFE class provides an interface for specifying a distribution of the fitness effects (DFE) of new mutations. Each DFE object specifies a list of mutation types and their proportions, where each mutation type is associated with a dominance coefficient, the distribution of selection coefficients (e.g., fixed-value, normally-distributed, gamma-distributed), and the parameters of this distribution. DFE objects can also model a relationship between the selection and dominance coefficients (Huber, Durvasula, et al., 2018, see **Methods**). Each Annotation object represents a collection of functional genomic elements (e.g., exons, introns, intergenic regions, etc.) whose coordinates are specified using a GFF3 annotation file. More concretely, consider implementing a model of selection consistent with the DFE inferred by Kim et al. (2017) for exons in the human genome. The associated DFE object is defined with two mutation types, neutral and deleterious, corresponding to synonymous and non-synonymous sites, with their proportions set to their expected prevalence in exons. The neutral mutation type is associated with a fixed selection coefficient (0), and the deleterious mutation type is associated with a dominance coefficient of 0.5 and a gamma-distributed selection coefficient with mean and shape as inferred by Kim et al. (2017).

As with other aspects of stdpopsim, we intend the selection models to be as biologically accurate as possible. Thus, the stdpopsim catalog provides Annotation objects based on species’ publicly available functional genomic elements and DFE objects based on published DFE estimates (Figure 1A). By using annotation objects, one can, for example, simulate populations where selected mutations (whose selection coefficients are drawn from a specified DFE) occur within annotation regions (e.g. exons), while all mutations occurring outside of these regions do not affect fitness. There are relatively few species with published DFE estimates in the literature. stdpopsim currently has implemented DFEs for five species in the catalog: *Arabidopsis thaliana* (Huber, Durvasula, et al., 2018), *Drosophila melanogaster* (Ragsdale, Coffman, et al., 2016; Huber, Kim, et al., 2017; Zhen et al., 2021), humans (Huber, Kim, et al., 2017; Kim, Huber, and Lohmueller, 2017; Kyriazis, Robinson, and Lohmueller, 2023; Castellano et al., 2019; Castellano et al., 2019; Rodrigues, Kern, and Ralph, 2024), *Mus musculus* (Booker et al., 2021), and the vaquita porpoise *Phocoena sinus* (Robinson et al., 2022). While the DFEs are from a limited number of species, these DFEs can be applied to other species if and when it is suitable, namely, when a species of interest lacks a published DFE in the catalog. For instance, one could simulate a butterfly species with a gamma-distributed DFE originally estimated from humans, or a human population with a DFE estimated from *Drosophila*. Indeed, the DFE labeled HomSap/Mixed K23 is recommended by Kyriazis, Robinson, and Lohmueller (2023) for “general” use across mammals, when lacking a more specific DFE. Furthermore, the user can specify a custom DFE and provide their own annotations of functional elements to simulate selection in a species for which we do not yet have a published DFE included in the catalog.

**Figure 1:**
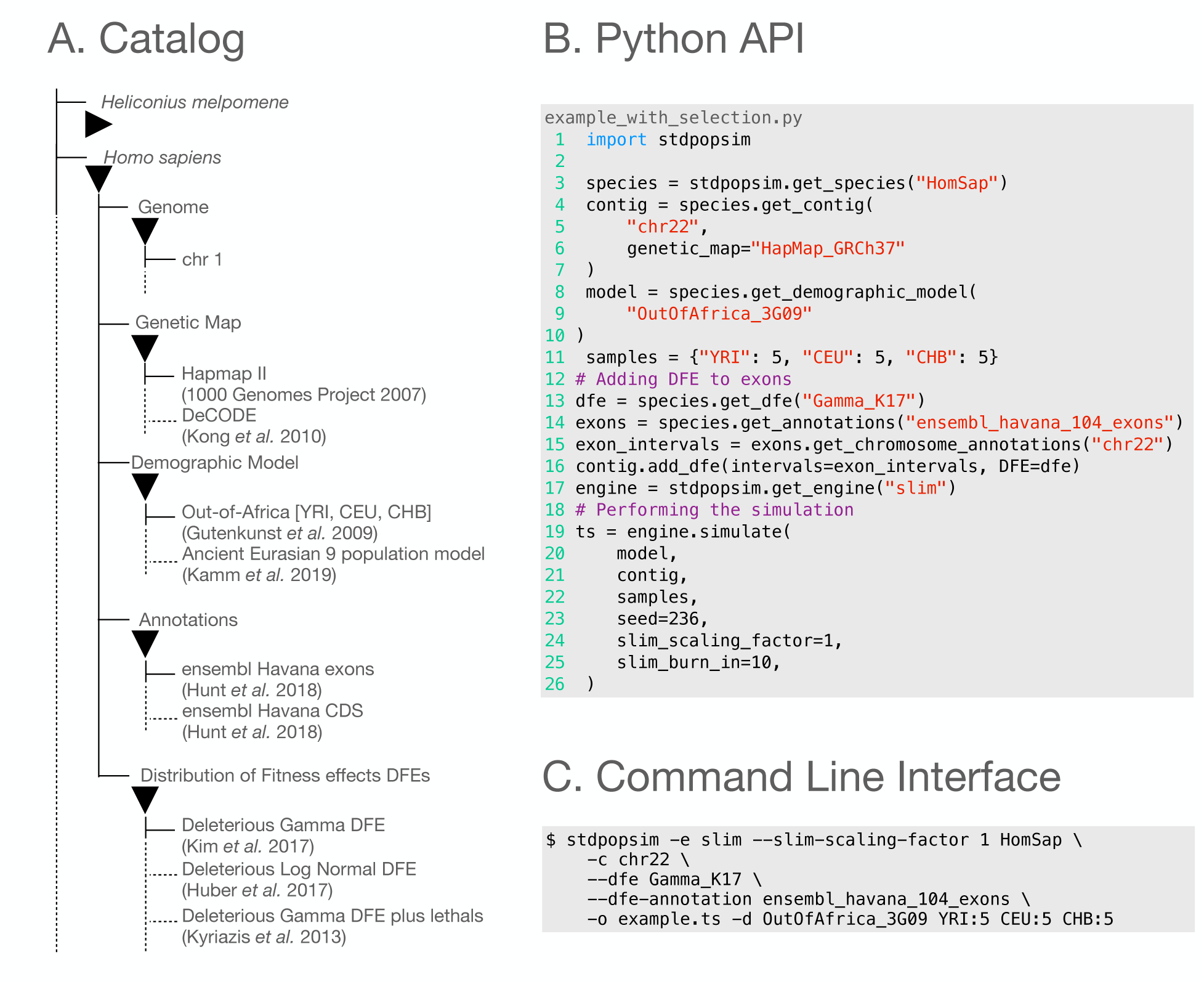
**(A)** A schematic of the stdpopsim catalog and the different components that can be used in simulations. **(B)** The Python API and **(C)** the command line interface. The user specifies a species, a portion of the genome to simulate, and optionally a genetic map, a model of demography, and a model of selection that itself is composed of a distribution of fitness effects (DFE) and a set of functional annotations.

From a user’s perspective, a model of selection is specified by pairing a DFE object with a collection of genomic segments defined using an Annotation object. Multiple such pairings can be defined in one simulation to provide a rich model of how selection may vary along a chromosome. For example, coding sequences may be associated with one DFE and non-coding sequences in exons with another DFE. We note that if a genomic interval belongs to more than one Annotation object, then it will be eventually associated with the last Annotation object added to the simulated contig and removed from any previously-added annotations. Simulations with selection are then implemented by specifying a species, a portion of the genome to simulate, a genetic map if available, a model of demography, and pairings of DFEs with genomic segments using either the Python API or the command-line interface (Figure 1B-C). Simulations with selection are generated using the SLiM simulation engine, which is perhaps the most flexible simulation engine for modeling selection available to date (msprime’s selection models are relatively limited and thus not currently used by stdpopsim).

While the DFE class facilitates the simulation of non-neutral variation across a sequence, it does not provide a means of specifying the origin and fate of particular selected alleles. Thus, we have also implemented an API in stdpopsim (stdpopsim.selective sweep) that enables selective sweeps to be simulated with a high degree of control. This function augments a demographic model by introducing a mutation at a particular time and position along the sequence; sets the selection coefficient of the mutation to a positive value over a given time interval; and conditions the simulation such that the mutation is above a particular frequency at the beginning and end of the sweep period. The conditioning is implemented via rejection sampling; if the advantageous allele is lost or does not have a frequency exceeding a user-specified threshold at the beginning or end of the sweep, the simulation is restarted from the introduction of the advantageous allele. If there are competing advantageous alleles (e.g. simultaneous sweeps at distinct loci) then the simulation is restarted from the introduction of the first of these (see **Methods**). As this functionality is implemented on top of the existing stdpopsim API, it may be combined with other models of selection and demography to simulate the combined effects of multiple, disparate evolutionary processes on genetic variation in a population.

At this release (version 0.3.0) the stdpopsim catalog comprises 27 species (i.e. genome representations), including vertebrates, invertebrates, plants, and bacteria, for which we have included 35 demographic models, 38 genetic maps, 12 DFEs, and 8 annotation objects. Additions to the catalog are ongoing—we point the reader to our previously published report detailing this effort (Lauterbur et al., 2023). We welcome contributions from the community and encourage those interested in contributing to visit https://popsim-consortium.github.io/stdpopsim-docs or contact one of the PopSim Consortium members for guidance.

### Inference of *N_e_*(*t*) in the context of selection

One of the most common applications of population genetic inference is to estimate the effective population size over time, *N_e_*(*t*), from genetic data. This can be done using a variety of methods, including the sequential Markovian coalescent (Li and Durbin, 2011; Schiffels and Wang, 2020; Terhorst, Kamm, and Song, 2017), through use of the site frequency spectrum (SFS) (Liu and Fu, 2020), as well as through linkage-disequilibrium information (Santiago et al., 2020). Since all of these methods assume neutrality, they are typically applied to genomic segments that are not expected to be directly affected by selection (e.g., by masking exons and conserved elements). However, selection acting on linked sites—even those that are masked out of the analysis—can bias estimates of *N_e_*(*t*) away from census population sizes by increasing the rate of coalescence in the genome (e.g. Schrider, Shanku, and Kern, 2016). Using stdpopsim, we can readily examine how different methods for inferring *N_e_*(*t*) are influenced by selection on linked sites, which in the case of negative selection is commonly referred to as *background selection* (Charlesworth, Morgan, and Charlesworth, 1993; Hudson and Kaplan, 1995).

To do this we simulated human genomes with and without selection. In both scenarios, we ran three replicate simulations under the out of Africa (OOA) demographic model of Ragsdale and Gravel (2019) using a genetic map from the HapMap Project (International HapMap Consortium et al., 2007) (see **Methods**). Simulations with selection were implemented by modeling purifying selection on nonsynonymous mutations using a DFE inferred by Kim, Huber, and Lohmueller (2017). A basic summary of nucleotide diversity from these simulations shows that the presence of selection decreases polymorphism only slightly at the genome-wide level, but more substantially within exonic regions (Figure S1).

Following the simulation, inference of *N_e_*(*t*) was conducted from each of the six simulated datasets using four methods: stairwayplot (Liu and Fu, 2020), msmc2 (Schiffels and Wang, 2020), SMC++ (Terhorst, Kamm, and Song, 2017) and GONE (Santiago et al., 2020). Although the estimates produced by the four methods differ from one another, for each method the estimates produced from data with selection (Figure 2; orange) and without selection (Figure 2; blue) are fairly similar. Thus, this particular parameterization of background selection and demography, in combination with selection only at exonic sites, does not appear to notably bias the estimates produced by these methods. This makes sense in light of our observation that genetic diversity is mostly affected within exonic regions and presumably sites very closely linked (Figure S1), thus masking of exons should be sufficient to remove the major effects of selection. As noted in previous studies that compared demographic inference methods on neutral simulations (Adrion et al., 2020), we see that each method has its strengths and weaknesses. Methods based on the sequential Markovian coalescent (SMC++ and msmc2) appear to produce the most accurate trajectories of *N_e_*(*t*) overall in this setting, although they tend to under-estimate the population bottleneck associated with the population split. On the other hand, stairwayplot, which utilizes the site frequency spectrum, produces more noisy estimates, but it does not appear to under-estimate the population bottleneck. The inference of GONE, which utilizes linkage-disequilibrium, is targeted toward demographic changes in more recent time frames (∼ 200 generations), explaining its noisy estimates for all but the most recent time periods.

**Figure 2:**
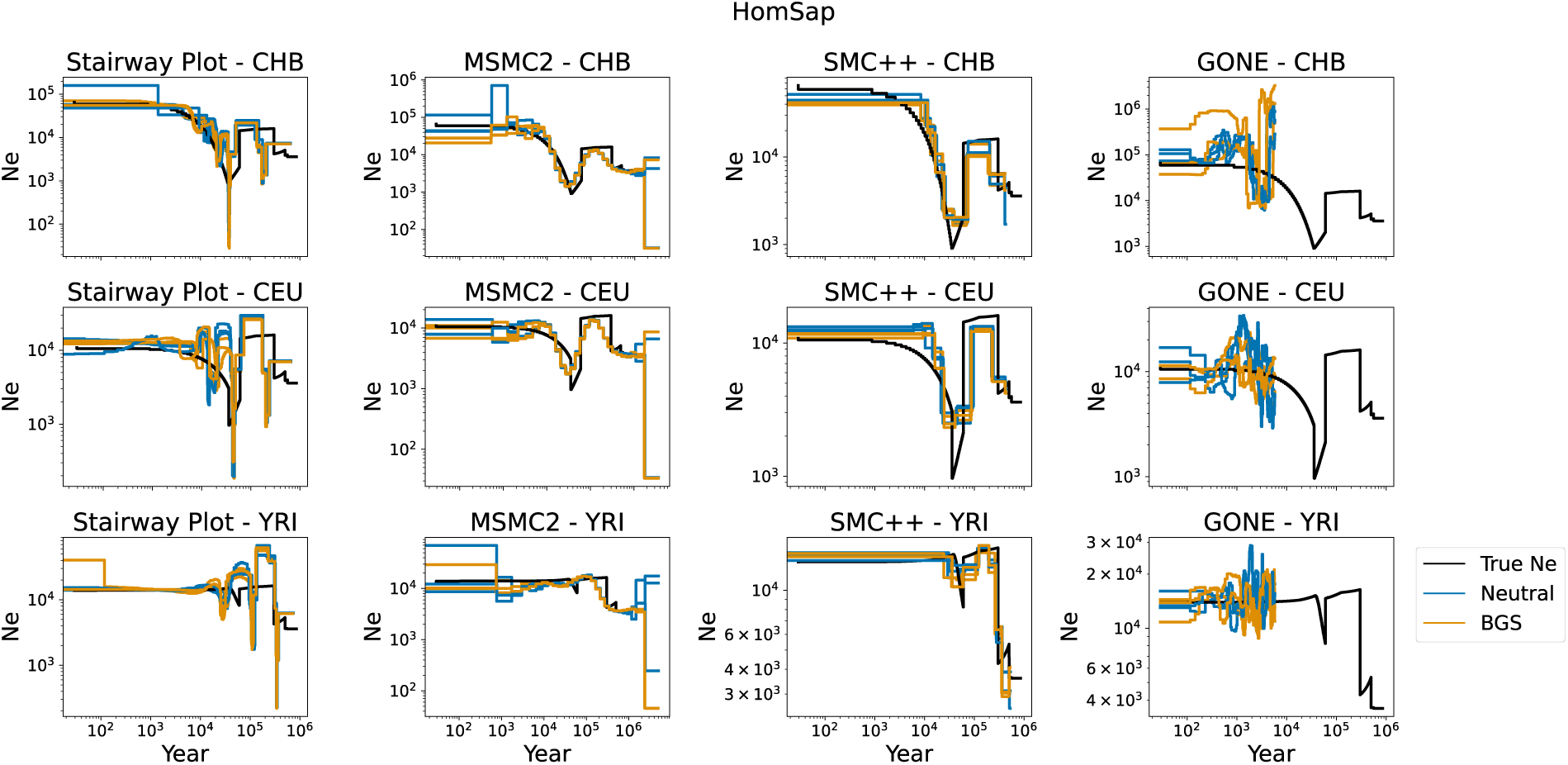
Inference of *N_e_*(*t*) from human genomes simulated under an out-of-Africa demographic model with and without purifying selection on exons. Rows correspond to the three extant populations in the simulation: CHB, CEU, and YRI. Columns correspond to the four inference methods: stairwayplot, msmc2, SMC++ and GONE. Each plot depicts the inferred *N_e_*(*t*) on the three datasets simulated without selection (blue) and the three datasets simulated with the background effects of purifying selection on exons (orange), alongside the true values of *N_e_* used in simulation (black). The X axis depicts time back from present (in years) and the Y axis depicts *N_e_* (number of individuals).

To expand these observations to additional species and to highlight the ease of comparison enabled by stdpopsim, we performed a similar analysis using the vaquita porpoise *Phocoena sinus*. The vaquita offers an intriguing contast to the demographic history of humans. While human poulations continue to expand, the vaquita is critically endangered, with only approxmiately 10 individauls remaining. Previous demographic inference from genetic data Robinson et al. (2022) found a long-term effective population size of 4,485 indiviuals which then decreased to 2,807 individuals 2,162 generations ago. The population then severely decreased even more to its current size in the last few generations. We simulated 100 genomes under a two-epoch demographic model inferred for vaquita by Robinson et al. (2022), and a constant recombination rate across the genome (see **Methods**). This is particularly interesting because the model was inferred assuming neutrality (with dadi) and used to inform conservation efforts for this critically endangered species. In simulations with selection, we applied the DFE inferred by Robinson et al. (2022) to exons annotated from the vaquita genome assembly. This DFE implements a relationship between selection and dominance in which very deleterious mutations (with selection coefficient *s* < −0.1) are fully recessive (dominance coefficient *h* = 0) and weakly deleterious mutations (*s* ≥ −0.001) are nearly additive (*h* = 0.4; see **Methods**). As in the human simulations, we see that the presence of selection decreases polymorphism in genomes simulated with selection, but that the effect is only readily apparent within exonic regions (Figure S2). For these simulations, the SFS-based method (stairwayplot) produces more accurate inference than the two methods based on the sequential Markovian coalescent (SMC++ and msmc2; see Figure S3). Regardless of these differences, we see that background selection again does not appear to considerably influence the inference of any of the four methods for this species, at least for the parameterization used here, even though vaquita has a very different evolutionary history than humans.

### Estimation of the Distribution of Fitness effects

Another common application of population genetic inference is to estimate the distribution of fitness effects (DFE) of new mutations from genetic data. The new framework of modeling selection in stdpopsim makes it an ideal tool for easily benchmarking and comparing the performance of different methods for inferring the DFE. Here, we considered three DFE-inference methods: GRAPES (Galtier, 2016), polyDFE (Tataru and Bataillon, 2020), and dadi-cli (Huang et al., 2023).

A challenge when comparing different inferred and simulated distributions of the selection coefficient (*s*) is differing conventions in the literature regarding its definition. The simulator SLiM and the inference tool GRAPES define the selection coefficient *s* such that a homozygote has fitness 1+*s*. On the other hand, dadi-cli and polyDFE define the selection coefficient such that a homozygote has fitness 1 + 2*s*. Moreover, inferred distributions of selection coefficients are typically scaled by an inferred ancestral effective population size *N_e_*, and different methods assume different scaling. dadi-cli infers the distribution of 2*N_e_s* while polyDFE and GRAPES infer the distribution of 4*N_e_s*. To allow direct comparison, here we have normalized all inference results to the convention used by SLiM (also used by stdpopsim; see **Methods**).

We start by examining DFE inference from human genomes simulated under an out-of-Africa demographic model with selection acting on exons (see **Methods**). In this setting, the DFE is inferred separately for each of the three simulated populations using each of the three DFE-inference methods. In each case we note that the DFE is estimated from segregating sites only, without the use of substitution data, as stdpopsim’s models currently do not contain outgroup populations or species that can be used to estimate divergence.

Overall, inferred DFEs varied across populations and inference methods (Figure 3, Table S1). Note that while the estimation accuracy for the DFE parameters themselves is informative (Figure 3A-B), examining the difference between the actual and inferred distributions of *s* can be more helpful for showing how well a method has recovered the true DFE (Figure 3C). The distributions of *s* inferred by polyDFE and dadi-cli had higher mean values (in absolute value) but smaller shape parameters, while GRAPES inferred distributions with lower mean values and shape parameters. As a result, polyDFE and dadi-cli over-estimate the fraction of mutations under strong selection (*s* < −0.1), while GRAPES considerably over-estimates the fraction of mutations under weak selection (*s* ≥ −0.0001).

**Figure 3:**
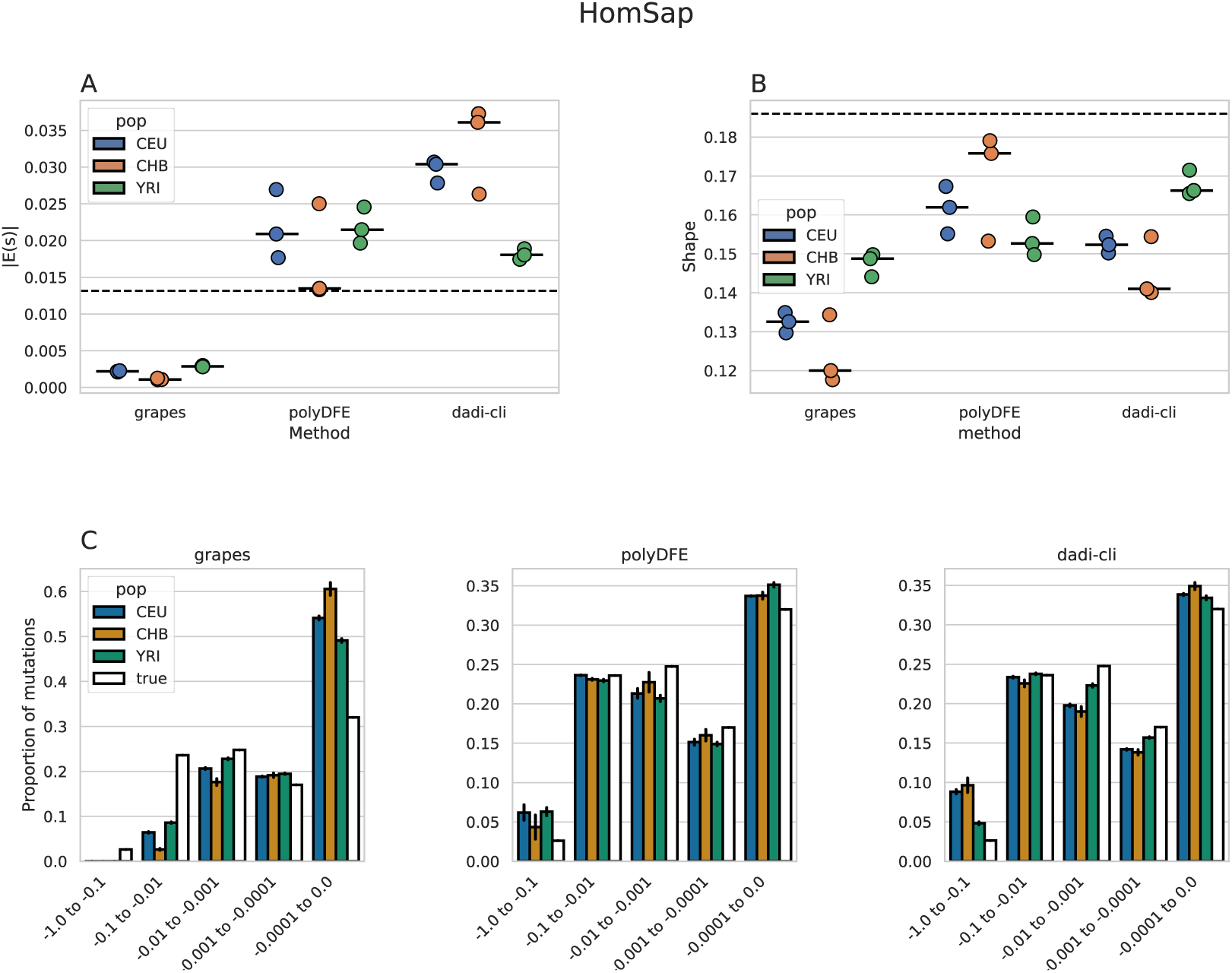
Inferred versus simulated DFEs from simulated human genomes. Simulations were performed using a human out-of-Africa demographic model with a gamma-distributed DFE acting on exons (see **Methods**). DFE is inferred separately for each of the three extant populations (CEU, CHB, and YRI) by three different methods: GRAPES, polyDFE, and dadi-cli. (A-B) Mean absolute value of selection coefficient (*|E*(*s*)|) and shape parameter are shown for each DFE inferred from all three simulation replicates, with median values marked by horizontal bars and simulated values represented by dashed horizontal lines. (C) Binned distribution of *s* implied by the average DFE inferred from the three simulated datasets (averaging the inferred gamma parameters), with error bars depicting the standard error of the mean for each bin. White bars represent the distribution used in the simulation.

Population history appears to influence different methods in slightly different ways. For instance, dadi-cli and GRAPES produce more accurate estimates from the YRI genomes (relative to the other two populations), whereas polyDFE shows very similar accuracy across all populations. To factor out the influence of demographic changes, we also analyzed genomes generated using similar simulations but with no population size changes (Figure S4, Table S1). In this simpler setting, polyDFE overlapped the true mean selection coefficient and shape parameter, whereas the other methods do not show as close a correspondence.

GRAPES exhibits similar biases as observed in the simulations with more complex demography, but with milder under-estimation of the shape parameter, whereas dadi-cli shows an inverse pattern with overestimated shape parameter and slightly underestimated mean. While providing possible insights into the effects of the demographic history on accuracy of different DFE-inference methods, substantiating these insights will require more than three replicate simulations per scenario.

Next, we examined the performance of these three DFE-inference methods when assuming a different DFE. To do this, we again turned to the vaquita. Unlike the human simulations where all mutations were assumed to be additive (h=0.5), the vaquita DFE applied a gamma-distributed selection coefficient to mutations in exons with a relationship between the selection and dominance coefficients (see **Methods**). Specifically, more deleterious mutations in the vaquita DFE are assumed to be more recessive. In this setting, all methods perform uniformly worse than in the human genome simulations, with consistent underestimation of the mean selection coefficient and overestimation of the shape parameter (Figure 4A-B, Table S1). Similar patterns are observed when re-running the simulations with a simpler demographic model without population size changes (Figure S5, Table S1). We suspect that this is due to the fact that all DFE-inference methods used here assume an additive model 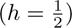, while mutations in the simulated genomes were more recessive 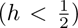, and their dominance decreased with the strength of selection. To examine the influence of this model misspecification, we compared the inferred distributions of *s* to the simulated distribution of 2*hs*, which would have been equal to *s* under an additive model (Figure 4C). Reassuringly, all three methods infer distributions that appear to fit the main mode of the simulated distribution of 2*hs*. This is likely because deleterious alleles are typically at low frequency and thus heterozygous, where their selective effect is *hs* rather than *s*.

**Figure 4:**
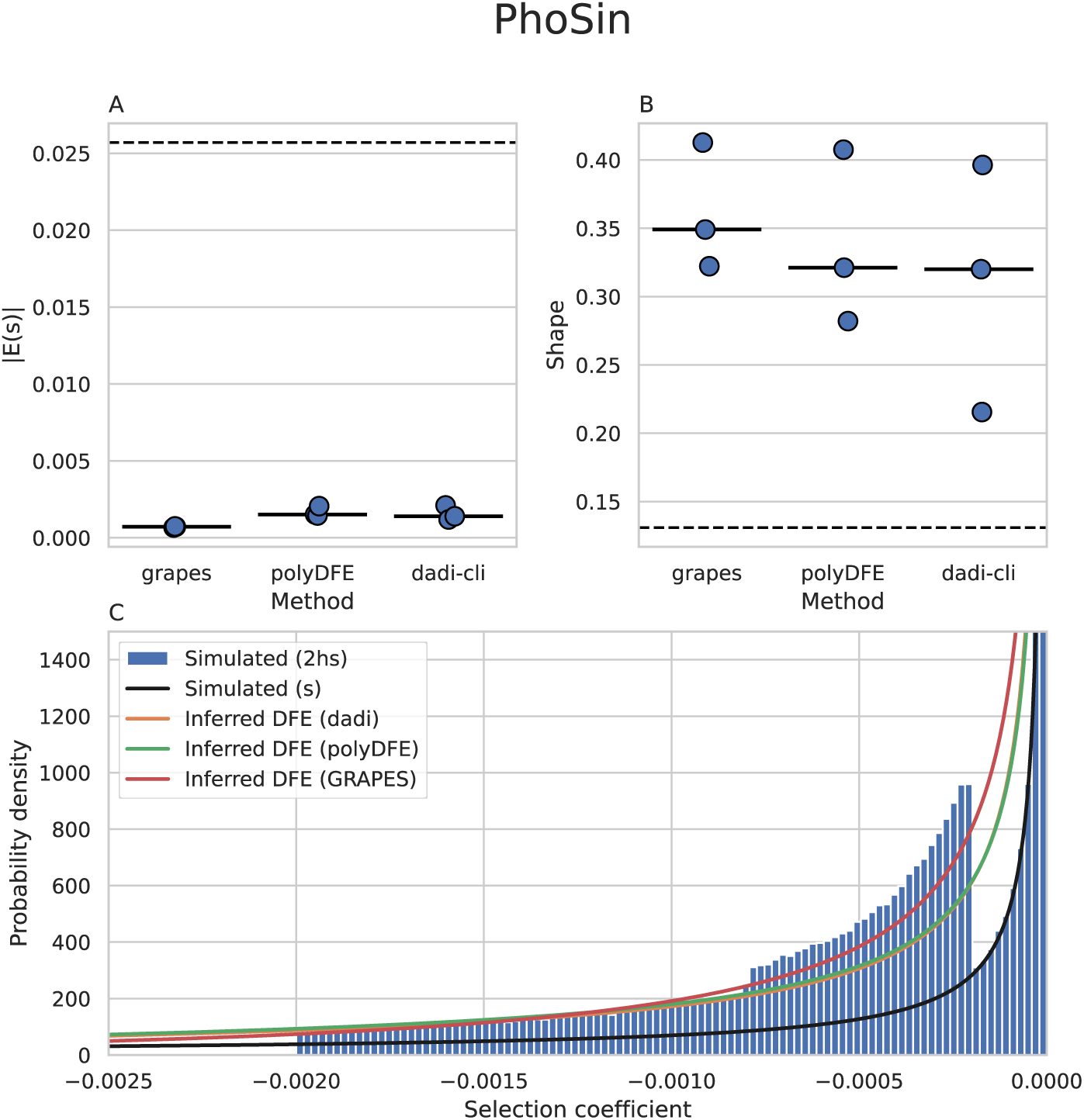
Inferred versus simulated DFEs from simulated vaquita genomes. Simulations were performed using a two-epoch model of vaquita porpoise demography with a gamma-distributed DFE acting on nonsynonymous mutations with a relationship between the selection coefficient (*s*) and dominance coefficient (*h*) (see **Methods**). The DFE is inferred by analyzing all simulated genomes jointly by one of three different methods: GRAPES, polyDFE, and dadi-cli. (A-B) Mean absolute value of selection coefficient (*|E*(*s*)|) and shape parameter are shown for each DFE inferred from all three simulation replicates, with median values marked by horizontal bars and simulated values represented by dashed horizontal lines. (C) Binned distribution of 2*hs* implied by the simulated DFE compared with the distribution of *s* implied by the average DFE inferred for each method from the three simulated replicates (averaging the inferred gamma parameters). The distribution of 2*hs* is multimodal because of the simulated relationship between *h* and *s* (see text).

### Detection of Selective Sweeps

We next highlight how stdpopsim can be used to benchmark and compare the performance of different methods for detecting selective sweeps from genetic data. Because simulation of selective sweeps is time consuming and evaluation of statistical power requires many replicate simulations, we focused on human chromosome 1 and generated replicate simulations on 5cM segments of the chromosome in 93 different locations. Selective sweeps were simulated by introducing a beneficial mutation in the center of the 5cM segment with a moderately strong selection coefficient (*s* = 0.03; 2*N_e_s* ∼ 600; see **Methods**). Each segment was simulated 1000 times, while keeping only replicates in which the selected allele reached a frequency of 95% or greater. These sweeps were modeled in the context of the three population out-of-Africa model from Gutenkunst et al. (2009) and background selection from deleterious mutations in exons. Sweep-detection methods typically base their inference on checking whether a test statistic exceeds a certain threshold, which is determined according to some empirical null distribution. In our simulations, we considered two different null distributions: one based on neutral simulations, and another based on simulations with background selection from deleterious mutations in exons (see **Methods**).

We evaluated the statistical power to detect selective sweeps for three methods: sweepfinder2 (DeGiorgio, Huber, et al., 2016), diploshic (Kern and Schrider, 2018), and reduced diversity (π). For each method, we determined a threshold for its test statistic using each of the two null models separately – neutral and with background selection (BGS) – and controlling for a 5% false positive rate. We then applied each method to data simulated with sweeps in one of two populations (YRI and CEU) in each of the simulated locations along human chromosome 1 (Figure 5). First and most strikingly, the power to detect sweeps varies considerably along the chromosome, and this variation is much greater than the variation in power between CEU and YRI or between null models. This is likely due to the effects of recombination rate heterogeneity (Figure 5). Next, we notice that power to detect sweeps is slightly lower when the test statistic threshold is determined using simulated data with background selection (BGS null model). This is likely because the strength of selection acting on exons is relatively weak, and the sweep detection methods we have used are relatively robust to the effects of background selection. Finally, the power to detect sweeps is generally higher in the YRI sample than in the CEU sample, which is consistent with the fact that YRI has a larger ancestral effective population size and thus both stronger selection and more variation to detect a sweep (e.g., Simonsen, Churchill, and Aquadro, 1995). Conversely, the CEU population experienced the out-of-Africa bottleneck that may mute the signal of a sweep.

**Figure 5:**
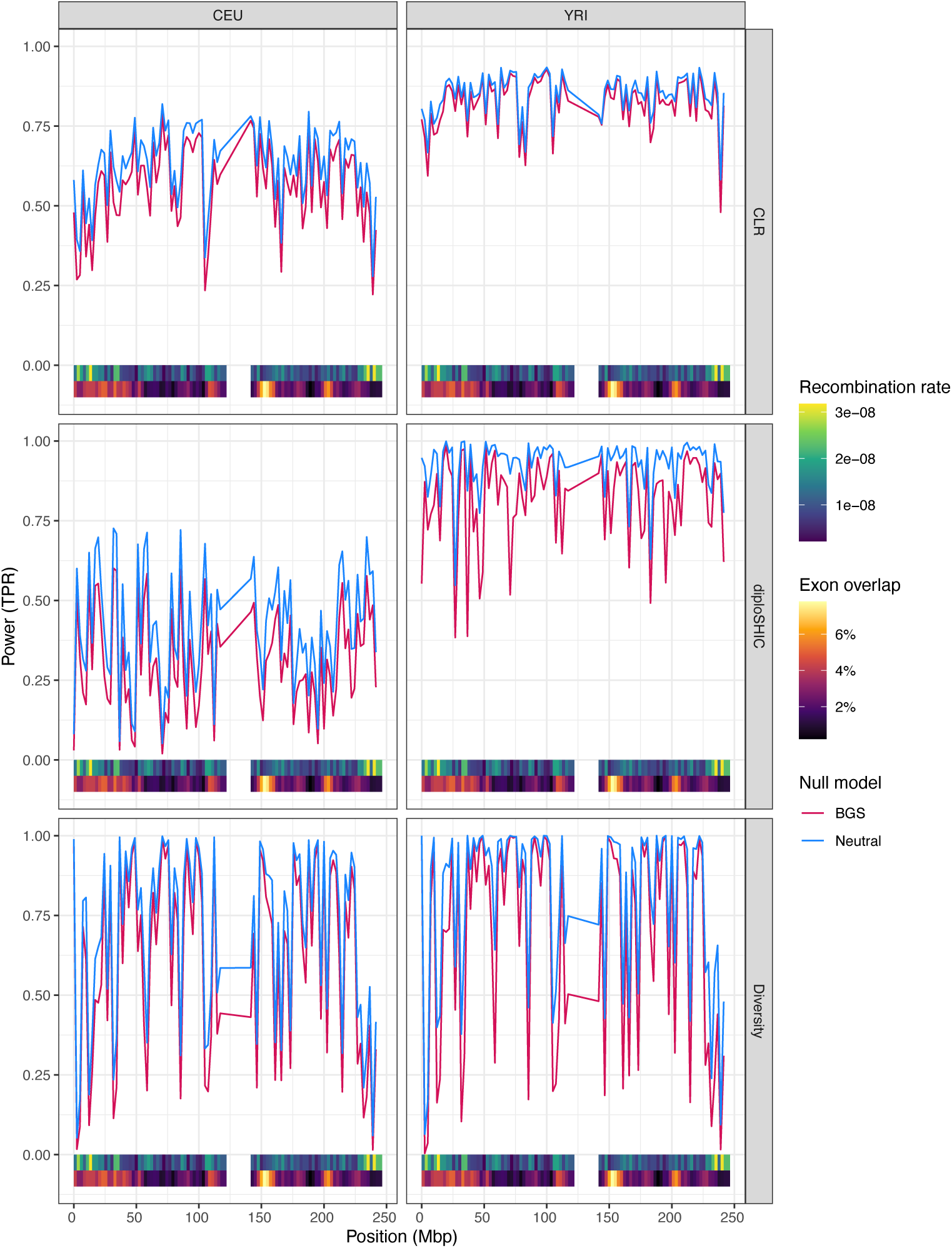
Power to detect selective sweeps along human chromosome 1. Genomic segments were simulated with sweeps under a three population out-of-Africa model and with background selection from deleterious mutations in exons. Three methods for detecting sweeps were applied to simulated data: sweepfinder2 (top row – labeled CLR), diploshic (middle row), and reduced diversity (π) (bottom row). Power (true positive rate) is shown for these methods for the CEU and YRI samples (left and right respectively). The thresholds of the test statistics were set to control for 5% false positive rate under a neutral null model (blue) and a null model with background selection from deleterious mutations in exons (red). Also shown are heatmaps of local recombination rate and exon density along the chromosome (bottom).

We compared the overall performance of the three sweep-detection methods using receiver operating characteristic (ROC) curves (Figure S6). Performance varies for each of the sampled populations. In the CEU sample, sweepfinder2 and diploshic perform similarly with a high area under the curve (AUC) score, (AUC = 0.906 and 0.903 respectively), but both underperform the simple method that uses reduced diversity (π) as an indicator (AUC = 0.932). In the YRI sample, diploshic outperforms the other two methods, with an AUC of 0.979, followed by using outliers of π with an AUC of 0.957, and then sweepfinder2 with an AUC of 0.939. We note that we trained diploshic on simulations under a constant size population, which more closely matches the demographic history of YRI, than that of CEU, which is characterized by a strong recent bottleneck followed by population expansion. Interestingly, the simple method that uses reduced diversity (π) as an indicator for a selective sweep performs very well overall (especially on the CEU sample). However, its accuracy is much more variable along the chromosome compared to the other two methods (see Figure 5), and for quite a few locations along the chromosome it has power below 50% when applied to both sample sets. sweepfinder2 appears to have the least variation in power along the chromosome or across populations.

To further dissect the reasons for high variance along the chromosome in power to detect sweeps, we examined the relationship between power and local recombination rate (Figure 6). We see that power to detect sweeps is a decreasing function of local recombination rate for sweepfinder2 and reduced diversity, which is consistent with sweeps having a larger genomic footprint in regions of low recombination. diploshic on the other hand shows a slight increase in power with recombination rates, presumably because it is designed to distinguish regions where a sweep happened from regions linked to a sweep, and if the recombination rate is too low (or selection too strong) then the genomic window examined by diploshic may not be large enough to exhibit enough of a recovery in patterns of diversity flanking the sweep. These results confirm that reduced genetic diversity performs well as an indicator for selective sweeps only in regions with low recombination rate. The other two methods, which were specifically designed for sweep detection, are considerably more robust to the effects of recombination rate than the simple indicator of reduced genetic diversity. These methods are particularly robust to recombination rate variation when applied to the YRI sample, especially when using a neutral null model to calibrate the test statistics. The other null model, which assumes background selection from exons, appears to consistently lead to reduced power for detecting sweeps regardless of the recombination rate. Interestingly, we do not see a clear correlation between power to detect sweeps and local exon density (Figure S7). This is somewhat surprising given that background selection may be expected to interfere with the sweep signature (reviewed by Stephan (2010)). This might suggest that the effects of realistic background selection on genome variation may not be as similar to those of sweeps as previously thought (Schrider, 2020).

**Figure 6:**
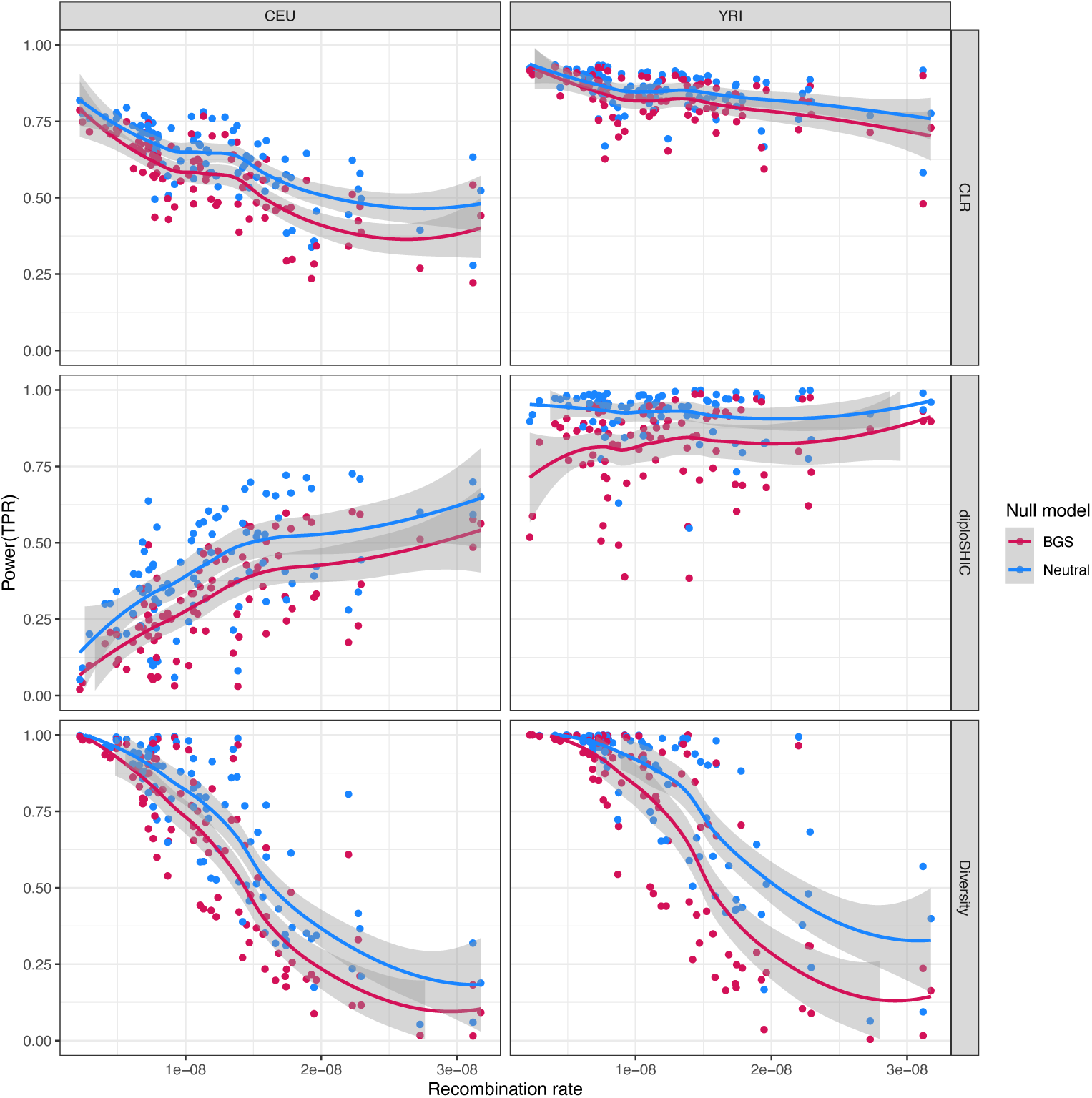
Power to detect selective sweeps as a function of local recombination rate. This figure shows the same power estimates shown in Figure 5, but with the genomic segments plotted against their average recombination rates instead of position along chromosome 1. Genomic segments were simulated with sweeps under a three population out-of-Africa model and with background selection from deleterious mutations in exons. Three methods for detecting sweeps were applied to simulated data: sweepfinder2 (top row– labeled CLR), diploshic (middle row), and reduced diversity (π) (bottom row). Power (true positive rate) is shown for these methods for the CEU and YRI samples (left and right respectively). The thresholds of the test statistics were set to obtain a 5% false positive rate under a neutral null model (blue) and a null model with background selection from deleterious mutations in exons (red). Fitted lines represent loess smoothed regressions.

## Discussion

In this paper, we have presented an important new addition to the stdpopsim library that enables the simulation of genetic variation in the presence of selection. We have demonstrated the utility of this new API by showing several illustrative examples of how it can be used to benchmark the performance of different methods for demographic history inference, the inference of the distribution of fitness effects of new mutations, and the detection of selective sweeps. This allows for a wide range of models of selection to be easily simulated and compared to empirical data, with reproducibility, computational efficiency, and rigor.

The exploration of performance among methods done here is not meant to be thorough benchmarking. Rather, we include these analyses as examples to highlight the utility of stdpopsim for both benchmarking and other tasks where researchers would benefit from the ability to perform simulations using parts of published population genetic models in a streamlined and quality-controlled fashion. Having said that, there are a number of interesting results that merit discussion. The first pertains to estimation of demographic history (*N_e_*(*t*)) in the context of linked purifying selection. Our simulations of human and vaquita genomes model deleterious mutations only at exonic positions of the genome. As shown in Figures S1 and S2, this combination only produces sizable decreases in polymorphism at exonic positions, which represent a small minority of sites within the genome, and does not lead to large-scale perturbations in genome-wide diversity. Thus, it is not surprising that estimates of *N_e_*(*t*) from genome-wide data from which exons have been masked should be accurate (e.g., Figures 2 and S3). Indeed, our results echo those of Marsh and Johri (2024) who found similar results on realistic human simulations that included selection. However, the question remains to what extent this would be true in empirical data where selection is not constrained to coding positions. For instance, we know that many genomes harbor large numbers of conserved non-coding elements (e.g., Siepel et al., 2005) with extremely strong selection acting on them (Katzman et al., 2007; McLean and Bejerano, 2008). We thus caution regarding extrapolation from our results. Future studies might build upon the annotations we used here to include selection at other portions of the genome as well as mixtures of DFEs across sites.

A second important finding is that inference of the DFE is sensitive to the dominance of deleterious mutations. All three of the DFE-inference methods applied here assume that deleterious mutations act in an additive fashion (*h* = 0.5) by default. However, the simulated DFE for the vaquita genome included highly recessive mutations. Specifically, these simulations used a DFE where mutations with *s* < −0.01 are quite recessive, with *h <*= 0.01, and where mutations with −0.01 < *s* < −0.001 have *h* = 0.1. We found that the three DFE-inference methods underestimated *E*[*s*] and overestimated the shape parameter (Figure 4A-B). This performance is not surprising since decreasing *h* for a given distribution of *s* will lead to a decrease in *hs*, or an apparent decrease in the amount of selection. Thus, fitting a mis-specified DFE that assumes *h* = 0.5 leads to underestimates of *E*[*s*]. Our analysis suggests that the distributions of *s* estimated by these methods provide a reasonable approximation for the true distribution of *hs* (Figure 4C). This points to a notable limitation of most existing DFE inference methods, which assume that fitness effects (*hs*) are gamma-distributed. As shown in Figure 4C, while a gamma distribution might fit the marginal distribution of *s*, a relationship between *h* and *s* might result in fitness effects (*hs*), which are not gamma-distributed.

Overall, these results are consistent with prior work showing that it is challenging to separately infer the DFE of *s* along with *h* (Wright (1937), Veeramah et al. (2014), Simons et al. (2014), Fuller et al. (2019), Balick et al. (2022), and Kyriazis and Lohmueller (2024); reviewed in Fuller et al. (2019)) and that a large proportion (*>* 5%) of recessive deleterious mutations can confound inferences of *s* (Wade et al., 2023). Given the plethora of evidence for recessive deleterious mutations (Mukai et al., 1972; Agrawal and Whitlock, 2011; Huber, Durvasula, et al., 2018; Di and Lohmueller, 2024), these results suggest the need for care in DFE inference. Kyriazis and Lohmueller (2024) inferred the DFE of *s* for nonsynonymous mutations in humans when considering different values of *h*. Intriguingly, additive models showed a satisfactory fit to the SFS of nonsynonymous variants in humans, and more complex models of dominance did not substantially improve the fit. Nevertheless, models including recessive deleterious mutations, like those simulated here for the vaquita genome, show a similar fit to the SFS and may be more biologically realistic than fully additive models, given the evidence for the presence of recessive deleterious mutations. More broadly, new computational approaches are needed to precisely quantify the DFE and the covariance of *h* and *s*.

Further, inference of the DFE was done here using exon annotations of the genome in question. A key distinction between our analyses and those done in empirical work is that we have complete knowledge of which sites are under selection in the simulation. Our inferences therefore represent a best case scenario. Empirical inference of the DFE is a much harder problem, since some mutations assumed to be free of selection might in reality be under selection, and the strength of selection can change over time or across environments.

Regarding sweep detection, the inclusion of selection models into stdpopsim allowed us to readily compare the effectiveness of different methods using data that reflects the demographic history, recombination rate variation, and genomic locations of negatively selected sites present in human populations. These results clearly demonstrate the impact of recombination rate variation, and underscore the importance of training methods on data with variable recombination rates (e.g., Schrider and Kern, 2017). In addition, our results illustrate a strength of likelihood methods like sweepfinder2 that do not require a predefined window size. Window-based methods, such as diploshic, tend to lag in performance when the genomic region affected by the sweep is larger than the chosen window size, as can occur with very strong sweeps or low levels of recombination. Moreover, while all the methods examined here retain adequate power for sweep identification across scenarios, including the inclusion of background selection, our results underscore the importance of developing methods with adequate statistical power even in the face of unknown demographic histories. Indeed, larger differences in power were observed when applying a method to genomes simulated with different demographic histories (i.e., YRI versus CEU) than when calibrating a method using a null model with and without background selection.

Although we have shown how the new stdpopsim API can be used to benchmark the performance of different methods, the comparisons shown here merely scratch the surface of what can be done with the new API. We envision that, moving forward, many researchers will use the new API to simulate data under a wide range of models of selection, demography, and genome architecture, and to develop new methods for the analysis of genetic data in the presence of selection. The additions described here further expand the utility of stdpopsim for both characterizing the patterns of diversity expected under a wide range of population genetic models, and for benchmarking methods for the analysis of genetic data on common ground in a way that was not possible before.

## Methods

### Annotations

Currently available annotation data in stdpopsim were downloaded from Ensembl. Coding sequence and exon annotations are available for *Arabidopsis thaliana* (Cheng et al., 2017), *Drosophila melanogaster* (Hoskins et al., 2015), *Homo sapiens* (Hunt et al., 2018a), and *Phocoena sinus* (vaquita porpoise, Morin et al., 2021).

### Implementing a DFE model using stdpopsim’s DFE class

The DFE class has the following attributes for modeling natural selection in a given set of annotated genomic segments. Similarly to other stdpopsim classes, the DFE class has fields for specifying the model identifier, description strings for documentation, and a citation object for the publication containing information on the inference on which the DFE model is based. A DFE model is specified using a mixture of mutation types, each associated with a frequency and a distribution of the seletion coefficient (*s*). Thus, each DFE object contains a list of MutationType objects and a separate list (of identical length) of positive frequencies (that sum up to 1). Each MutationType object is associated with a distribution type, distribution parameters, and dominance coefficients. The distribution types that are currently supported by stdpopsim are:

- Fixed (f), with one parameter specifying the fixed selection coefficient.
- Exponential (e), with one parameter specifying the mean of the exponential distribution.
- Gamma (g), with two parameters specifying the mean and shape of the gamma distribution.
- Normal (n), with two parameters specifying the mean and standard deviation of the normal distribution.
- Weibull (w), with two parameters specifying the scale and shape of the Weibull distribution.
- Uniform (u), with two parameters specifying the minimum and maximum of the interval.
- Positive log-normal (lp), with two parameters specifying the mean and standard deviation of the normal distribution in log scale.
- Negative log-normal (ln), with two parameters specifying the mean and standard deviation of the normal distribution in log scale.

Typically, one would associate a MutationType object with a single dominance coefficient (using the dominance coeff attribute): *h* = 0 for recessive mutations, 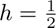 corresponding to an additive model (default value), and *h* = 1 for completely dominant mutations. The dominance coefficient, *h*, can take any real value, including negative values (for underdominance) and values greater than 1 (overdominance). There is also an option to model a relationship between the selection and dominance coefficients by associating different ranges of selection coefficients with different dominance coefficients. This is done using two attributes of the MutationType class: dominance coeff list is used to specify a list of *n* dominance coefficients, and dominance coeff breaks is used to specify *n* − 1 selection coefficient break points. The *i*th dominance coefficient in the list is associated with selection coefficients in the range between the *i* − 1th and the *i*th break points. For an explicit example, see the description of the vaquita porpoise genome simulations below. During simulation, when a mutation is generated in an annotated region associated with a given DFE object, a mutation type is sampled according to the frequencies of the different types. Then, a selection coefficient is sampled for that mutation based on the distribution associated with the sampled mutation type. Finally, a dominance coefficient is associated with the mutation either based on the fixed dominance coefficient of the mutation type (using attribute dominance coeff), or based on the sampled selection coefficient (using attributes dominance coeff list and dominance coeff breaks). For additional implementation details on the DFE class, see the stdpopsim docs at https://popsim-consortium.github.io/stdpopsim-docs/main/introduction.html.

### Selective sweeps and rejection sampling

Selective sweeps are implemented in the SLiM engine by introducing a mutation with a fitness effect of *hs* in heterozygotes and *s* in homozygotes, where *h* is the dominance coefficient (as for the DFE model) and *s* ≥ 0 is the selection coefficient. The allele is introduced in a given population at a specified time and genomic position, and may be either globally or locally adaptive (so that the fitness effect is confined to the population of origin) over a fixed duration. The start of the sweep may be offset from the introduction of the allele, to model selection on standing genetic variation (so-called “soft sweeps”).

Crucially, the advantageous allele must remain extant over the course of the sweep; this conditioning is accomplished via rejection sampling, whereby the simulation is restarted from the time of introduction of the mutation if the allele is lost in the population of origin. The conditioning may optionally be extended to reject trajectories where the frequency at the beginning and end of the sweep period is below a specified threshold. If there are competing advantageous alleles (simultaneous sweeps at distinct loci) and one of these does not satisfy the acceptance conditions, then the simulation is restarted from the onset of the first sweep; this avoids conditioning on particular (partial) allele frequency trajectories for the remaining alleles. A downside of this strategy is that rejection becomes more likely with increasing numbers of simultaneous sweeps or with hard-to-satisfy constraints; but it allows for greater flexibility in sweep specification and for interoperability with the stdpopsim DFE model.

### Quality control and software development

Following the protocol established previously for other aspects of stdpopsim (Adrion et al., 2020), new DFEs are subject to a rigorous quality control (QC) process, by which a different individual re-implements the same DFE, and the two implementations are programmatically tested for equality. This QC process has proven its effectiveness on numerous occasions in detecting implementation bugs and resolving modeling ambiguities. For example, this process detected an error in coding the log-normal distribution of the *Drosophila melanogaster* DFE inferred by Booker et al. (2021, stdpopsim label LognormalPlusPositive R16). This faulty DFE model was used in a study conducted by one of our groups before it underwent the QC process, where we noticed that deleterious selection coefficients were several orders of magnitude larger than expected. Subsequently, the QC process was able to find and fix this bug. In addition to QC, all changes and additions to the code must pass a comprehensive set of unit tests: at the time of writing, the library includes over 11,000 lines of testing code, comprising 2,862 unit tests, including comparisons to QC implementations and cross-checks for agreement, including between SLiM and msprime implementations of the same models. For instance, for testing we make use of SLiM’s scriptability to separately record as metadata the simulation parameters (current population sizes, DFEs, etc.), which are then programmatically compared to the models used to generate the SLiM script that ran the simulation. We make use of automatic test “code coverage” tools to help ensure that all portions of the code base are validated by these unit tests.

### Whole genome simulations

In this study we performed simulations of the human and vaquita porpoise genomes using stdpopsim. We ran replicate simulations for each genome/parameter combination, in which chromosomes were simulated separately from one another. All simulations were executed using the SLiM engine with a burn-in of 10*N* generations (where *N* is the initial population size), and a scaling factor of 1 (no scaling). We note that aggressive scaling of forward-in-time simulations can lead to quite different outcomes when selection is involved (Uricchio and Hernandez, 2014; Ferrari et al., 2024; Dabi and Schrider, 2025), and we caution users against using high degrees of scaling for drawing final inferences, although we recommend it during development to save on computational costs. All simulations were implemented using a snakemake workflow, where the code utilized is similar to that shown in Figure 1B. The complete simulation workflow is available at https://github.com/popsim-consortium/analysis2/blob/main/workflows/simulation.snake (see **Data availability** below).

### Simulations of complete human genomes

Simulation of human genomes used a genetic map from the HapMap Project (International HapMap Consortium et al., 2007, stdpopsim label HapMapII GRCh38) and the out of Africa demographic model with archaic admixture from Ragsdale and Gravel (2019, stdpopsim label OutOfAfricaArchaicAdmixture 5R19). Simulations with selection additionally applied a DFE model with gamma-distributed selection coefficients inferred by Kim, Huber, and Lohmueller (2017, stdpopsim label Gamma K17) to exons annotated by the HAVANA group release 104 (Hunt et al., 2018b, stdpopsim label ensembl havana 104 exons), which cover 2.1% of the genome. Each simulation generated 100 diploid genomes from each of three extant human populations (labeled YRI, CEU, and CHB), with each genome consisting of 22 autosomes. Using these settings, we generated three replicate datasets without selection and three replicate datasets with selection. Three additional datasets, used in Figure S4, were simulated with the same model of selection and a simple demographic model with one population with a constant size of 10, 000. Each of these simulations generated 100 diploid genomes. The configuration file used in combination with our snakemake workflow for these simulations can be found at https://github.com/popsim-consortium/analysis2/blob/main/workflows/config/snakemake/production_config_HomSap.yml. The running times of these simulations depend on chromosome length, the demographic model, and incorporation of selection (Figure S8).

### Simulations of complete vaquita porpoise genomes

Simulation of vaquita porpoise genomes used the two-epoch demographic model from Robinson et al. (2022, stdpopsim label Vaquita2Epoch 1R22). Since no genetic map is currently available for this species, we assumed a constant recombination rate of 10^−8^ across all chromosomes (Morin et al., 2021). Simulations with selection additionally applied a DFE model with gamma-distributed selection coefficients inferred by Robinson et al. (2022, stdpopsim label Gamma R22) to exons annotated from the vaquita porpoise genome assembly Morin et al. (2021, stdpopsim label Phocoena sinus.mPhoSin1.pri.110 exons), which cover 2.2% of the genome. This DFE model implements a relationship between selection and dominance as follows: very deleterious mutations (with *s* < −0.1) are recessive (*h* = 0), mutations with *s* ϵ [−0.1, −0.01) have a dominance coefficient of *h* = 0.01, mutations with *s* ϵ [−0.01, −0.001) have a dominance coefficient of *h* = 0.1, and very mild mutations (with *s* ϵ [−0.001, 0)) are nearly additive (*h* = 0.4). This discretized relationship matches that used by Robinson et al. (2022), and is specified using the following attributes in the DFE class: dominance_coeff_list= (0, 0.01, 0.1, 0.4), and dominance_coeff_breaks= (−0.1, −0.01, −0.001).

Each simulation generates 100 diploid genomes, with each genome consisting of 21 autosomes. Using these settings, we generated three replicate datasets without selection and three replicate datasets with selection. Three additional datasets, used in Figure S5, were simulated with the same model of selection and a simple demographic model with one population with a constant size of 3, 500. The configuration file used in combination with our snakemake workflow for these simulations can be found at https://github.com/popsim-consortium/analysis2/blob/main/workflows/config/snakemake/production_config_PhoSin.yml.

### Inference of *N*_e_(*t*)

Inference of *N_e_*(*t*) was conducted on simulated datasets after masking non-recombining portions of the genome (to avoid bias caused by extreme recombination rate variation) and exons (to avoid the direct effect of natural selection). Thus, inference from datasets simulated with selection was not influenced by direct selection acting on exons, but it was influenced by selection acting on linked sites (background selection). We assessed the accuracy of demographic inferences from four software packages: msmc2 (Schiffels and Wang, 2020), stairwayplot (Liu and Fu, 2020), GONE (Santiago et al., 2020), and SMC++ (Terhorst, Kamm, and Song, 2017). For msmc2 we used a random sample of *N* = 6 simulated genomes with 20 iterations of the EM algorithm. For GONE we set max snps to 500,000, the number of generations to 2,000, and the number of bins to 400, all representing default settings. For stairwayplot we did one run of *ε* estimation for each replicate dataset, and used a dimFactor of 5000, both of which represent default settings. For SMC++ we used the unfolded SFS (--unfold), along with the true mutation rate under which the simulations were generated. The complete snakemake workflow for demographic inference can be found at https://github.com/popsim-consortium/analysis2/blob/main/workflows/n_t.snake.

### DFE inference

We used three software packages to infer the distribution of fitness effects (DFE) from genetic data: dadi-cli (Huang et al., 2023; Kim, Huber, and Lohmueller, 2017), polyDFE (Tataru and Bataillon, 2020), and GRAPES (Galtier, 2016). Each software package was run using the same set of simulations. DFE inference using dadi-cli was conducted by first inferring a demographic model from neutral sites, using a two epoch demographic model with 100 repeated runs of the optimization procedure. dadi-cli then proceeds by using a cached strategy wherein we used 2000 points in the gamma distribution. Finally, the DFE was inferred using a ratio of non-synonymous to synonymous mutations of 2.31, again with 100 repeated runs of the optimization procedure. For polyDFE, we used model ‘C’ which uses a mixture of a gamma distribution and an exponential distribution, where the gamma component is used for negatively selected mutations. For optimization we used the default BFGS optimizer with the -e flag to automatically estimate the initial parameters values. For GRAPES, we inferred the DFE under the ‘GammaZero’ model, which assumes that selection coefficients are drawn from a single gamma distribution. We used the -no-div-param flag to disable the use of divergence-based parameter estimation. All other settings were left at their default values. All methods used the same exon annotations as generated in the simulations. The total number of coding sites in the human and vaquita genomes used were 61,644,801 and 31,039,513 respectively.

To facilitate direct comparison between inference methods, we scale each program’s output to obtain a selection coefficient (*s*) consistent with the one used by SLiM. GRAPES outputs the mean and shape of the inferred distribution of 4*N_e_s*. We thus scale the mean by a factor of 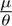, where *µ* is the assumed mutation rate and *ε* = 4*N_e_µ* is the population-scaled mutation rate, as estimated by GRAPES. polyDFE is similarly scaled by 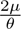, reflecting the fact that it defines the selection coefficient such that a homozygote has fitness 1 + 2*s*. dadi-cli defines the selection coefficient similarly to polyDFE and infers a gamma distribution for 2*N_e_s*, parameterized by the shape (*α*) and rate (*v*). It also infers the total population-scaled mutation rate for selected sites over the considered region *ε* = 4*N_e_µLp*, where *L* is the total number of sites available for the analysis, and *p* is the proportion of sites under selection. Thus, the mean value of *s* in SLiM terms is obtained by 4*µαvLp/θ*. Complete details for the DFE inference analysis can be found in the snakemake workflow at https://github.com/popsim-consortium/analysis2/blob/main/workflows/dfe.snake.

### Simulations with selective sweeps

To test methods for detecting selective sweeps, we considered 100 evenly distributed 5Mb segments of human chromosome 1. We included a 2.5cM buffer on either side of each segment to minimize the effect of chromosome ends on patterns of genetic variation. Seven segments in which the central 1Mb had an average recombination rate below 10^−11^ were excluded from the analysis. Each segment was simulated in each of the following three settings: (1) neutral, (2) background selection (BGS), and (3) hard sweep with BGS. In all three settings, we applied a genetic map from the HapMap Project (International HapMap Consortium et al., 2007, stdpopsim label HapMapII GRCh38) and the three-population out of Africa (OOA) demographic model from Gutenkunst et al. (2009, stdpopsim label OutOfAfrica 3G09). In the two settings with BGS, we applied the DFE model with gamma-distributed selection coefficients inferred by Kim, Huber, and Lohmueller (2017, stdpopsim label Gamma K17) to exons annotated by the HAVANA group release 104 (Hunt et al., 2018b, stdpopsim label ensembl havana 104 exons). Hard sweeps were modeled by generating a single beneficial mutation with selection coefficient of *s* = 0.03 in the center of the 5cM genomic segment and discarding simulations in which the beneficial mutation did not reach (or exceed) a terminal frequency of 95%. For each of the 93 genomic segments of chromosome 1, we generated 1,000 replicate datasets under each of the three settings. We used the SLiM engine with a burn-in of 2*N* generations (where *N* is the initial population size), and a scaling factor of 2. Each simulation generated ten diploid genomic segments for each of the two sampled populations: YRI and CEU. The sequences simulated with a hard sweep and BGS were used for sweep detection, and sequences simulated under the other two settings were used to calibrate the thresholds of the test statistics used by the three sweep-detection methods (see below). See Figure S9 for the distribution of runtimes for these simulations.

### Selective sweeps detection

Each of the three sweep-detection methods is based on a test statistic. Reduced nucleotide diversity is based on nucleotide diversity (π) computed directly from the underlying tree sequences using tskit (Ralph, Thornton, and Kelleher, 2020) over ten equally sized and non-overlapping subwindows. sweepfinder2 (DeGiorgio, Huber, et al., 2016) is based on a composite likelihood ratio (CLR) computed at 21 uniformly distributed positions along the simulated region. diploshic (Kern and Schrider, 2018) uses supervised learning to train a sweep classifier. Its training phase used discoal (Kern and Schrider, 2016) to simulate 20 haploid sequences of length 1.21Mbp with a constant population size of *N_e_* = 10,000, a mutation rate of 2 × 10^−8^, and a recombination rate of 2 × 10^−8^. For discoal simulations that included selective sweeps, selection coefficients were drawn from a uniform distribution between 0.001 and 0.10, and the time of the sweep was chosen from a uniform distribution between 0 and 0.01*N_e_* generations in the past. For soft sweeps, the frequency at which selection was introduced was drawn from a uniform distribution between 0.0001 and 0.1. We note that since the training set of diploshic was generated under a simple model with constant mutation and recombination rates and constant effective population size, we expect it to perform suboptimally on the simulated segments of human chromosome 1, which were generated using more realistic simulations (see **Simulations with selective sweeps** above).

The three sweep-detection methods were associated with thresholds for their respective test statistics (π, sweepfinder2’s CLR, or diploshic’s probability), which were determined using an empirical distribution representing a null model without sweeps. We considered two null models: a neutral evolution model (neutral), and a model with background selection from deleterious mutations in exons (BGS). Each null model was represented by a different set of 93, 000 simulations generated according to the appropriate model (see **Simulations with selective sweeps** above). Thus, for each test statistic we determined two thresholds corresponding to these two different null models (neutral and BGS) using the extreme 5% quantile in each empirical distribution. Then, power (or true positive rate) to detect a sweep by a given method was determined for positions on chromosome 1 as follows: each sweep-detection method was applied to the 1,000 simulated datasets generated with a hard sweep and BGS for that position, and the power was set to be the proportion of simulations (out of 1,000) in which the test statistic exceeded the threshold determined by each of the two null models (neutral and BGS). We note that using an empirical distribution to determine the threshold for calling a selective sweep with diploshic deviates from the original implementation of the method, however for consistency among methods we chose to take this approach. The complete snakemake workflow for sweep detection (including simulations) can be found at https://github.com/popsim-consortium/analysis2/blob/main/workflows/sweep_simulate.snake.

## Data availability

The release of stdpopsim is available at https://github.com/popsim-consortium/stdpopsim. This paper coincides with version 0.3.0 of stdpopsim. The entire analysis pipeline associated with this paper is available as a snakemake workflow at https://github.com/popsim-consortium/analysis2.

## Acknowledgments

We thank Tom Booker for discussions and encouragement to incorporate realistic BGS models into stdpopsim. We thank the numerous users of the stdpopsim package, as well as workshop participants who provided feedback and suggestions for improvements.

