## Supplemental Figures for "Accessible, realistic genome simulation with selection using stdpopsim"

1 # Supplementary Material

Table S1: **Performance of DFE-inference methods in four simulation scenarios**. For each of the three methods, we report the mean absolute error (MAE) of the mean selection coefficient and shape of its inferred DFE. In each simulation scenario, the lowest MAEs are highlighted in bold.

| species ID | demography | MAE $E\|s\|$ dadi-cli | MAE $E\|s\|$ grapes | MAE $E\|s\|$ polyDFE | MAE shape dadi-cli | MAE shape grapes | MAE shape polyDFE |
|---|---|---|---|---|---|---|---|
| HomSap | Constant | **0.0012** | 0.0101 | 0.0023 | 0.030 | 0.0086 | **0.0068** |
| HomSap | OOA 5R19 | 0.014 | 0.0111 | **0.0072** | 0.031 | 0.051 | **0.024** |
| PhoSin | Constant | **0.024** | 0.025 | **0.024** | **0.23** | 0.25 | **0.23** |
| PhoSin | Vaquita2Epoch 1R22 | **0.024** | 0.025 | **0.024** | **0.18** | 0.23 | 0.21 |

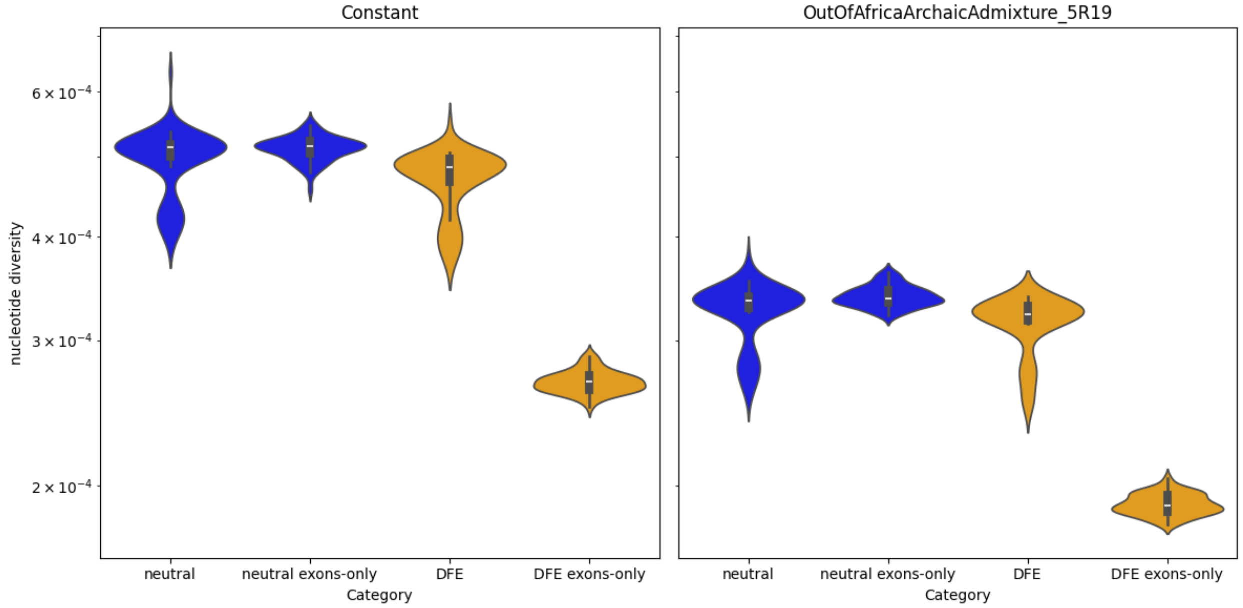


Figure S1: Genetic diversity in human simulations. Nucleotide diversity ($\pi$) was calculated for genomes simulated under constant population size (left panel) and under the out-of-Africa demographic model (right panel). Each panel shows four violin plots representing the distribution of $\pi$ in genomes simulated without selection (blue) and genomes simulated with background selection (orange), calculated from either the entire genome (labeled neutral or DFE), or from exonic regions only (labeled exons-only). Here, $\pi$ is computed across all simulated genomes (from the three simulated populations) considered as a single population. Each violin plot summarizes three replicate simulations generated under the relevant scenario.

2

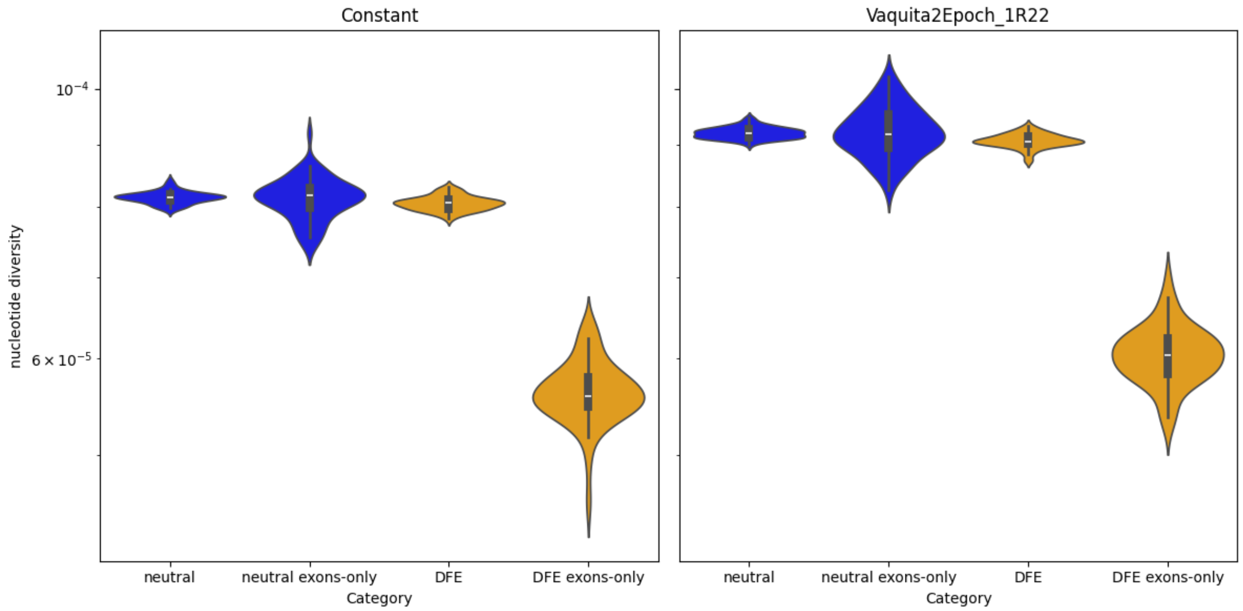


Figure S2: Genetic diversity in vaquita simulations. Nucleotide diversity ($\pi$) was calculated for genomes simulated under constant population size (left panel) and under the two-epoch demographic model (right panel). Each panel shows four violin plots representing the distribution of $\pi$ in genomes simulated without selection (blue) and genomes simulated with background selection (orange), calculated from either the entire genome (labeled neutral or DFE), or from exonic regions only (labeled exons-only). Each violin plot summarizes three replicate simulations generated under the relevant scenario.

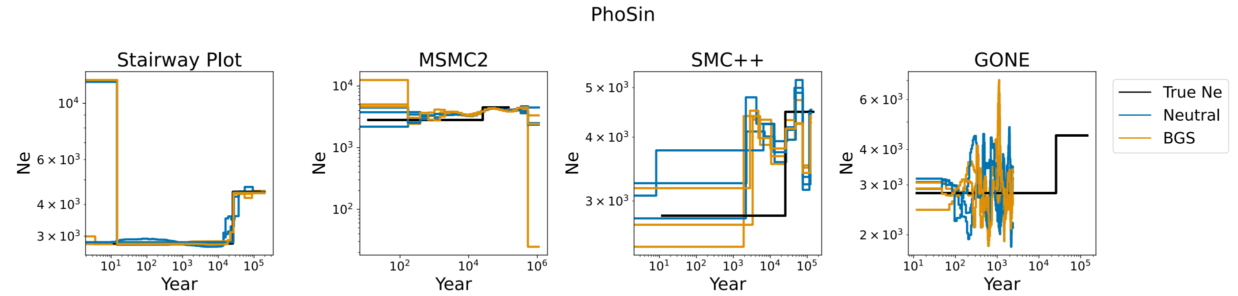


Figure S3: Inference of $N_e(t)$ from vaquita porpoise genomes simulated under a single population model of declining population size with and without purifying selection on exons. Inference is conducted using four methods: `stairwayplot`, `msmc2`, `SMC++` and `GONE`. Each plot depicts the inferred $N_e(t)$ on the three datasets simulated without selection (blue) and the three datasets simulated with the background effects of purifying selection on exons (orange), alongside the true values of $N_e$ used in simulation (black). The X axis depicts time back from present (in years) and the Y axis depicts $N_e$ (number of individuals).

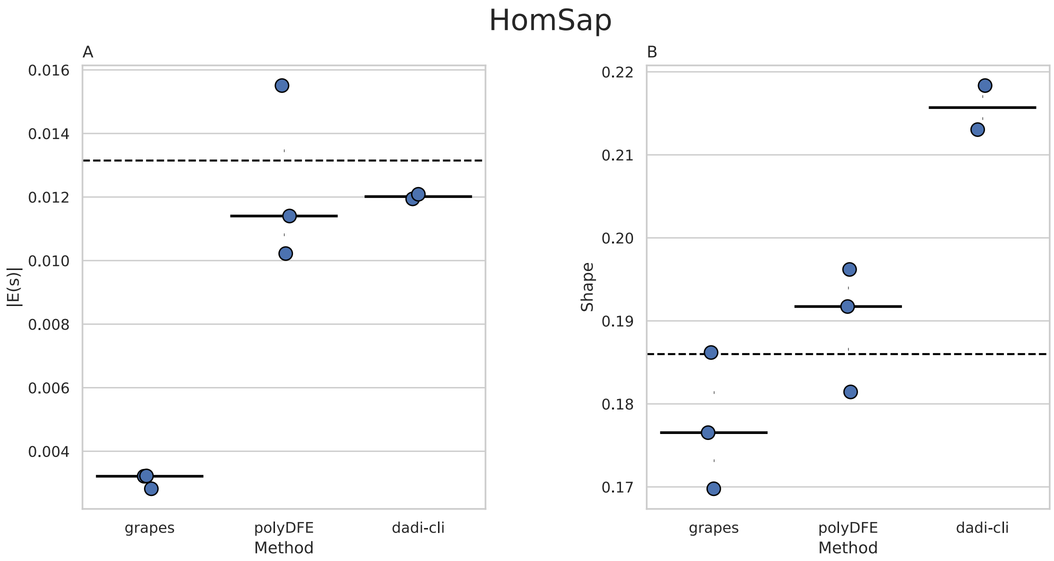


Figure S4: Inferred versus simulated DFEs from human genomes simulated with a simple demographic model with one population with a constant size of $10,000$ and a gamma-distributed DFE acting on exons (see **Methods**). DFE is inferred by three different methods: `GRAPES`, `polyDFE`, and `dadi-cli`. Mean absolute value of selection coefficient ($|E(s)|$, panel A) and shape parameter (panel B) are shown for each DFE inferred from all three simulation replicates with median values marked by horizontal bars and simulated values represented by dashed horizontal lines.

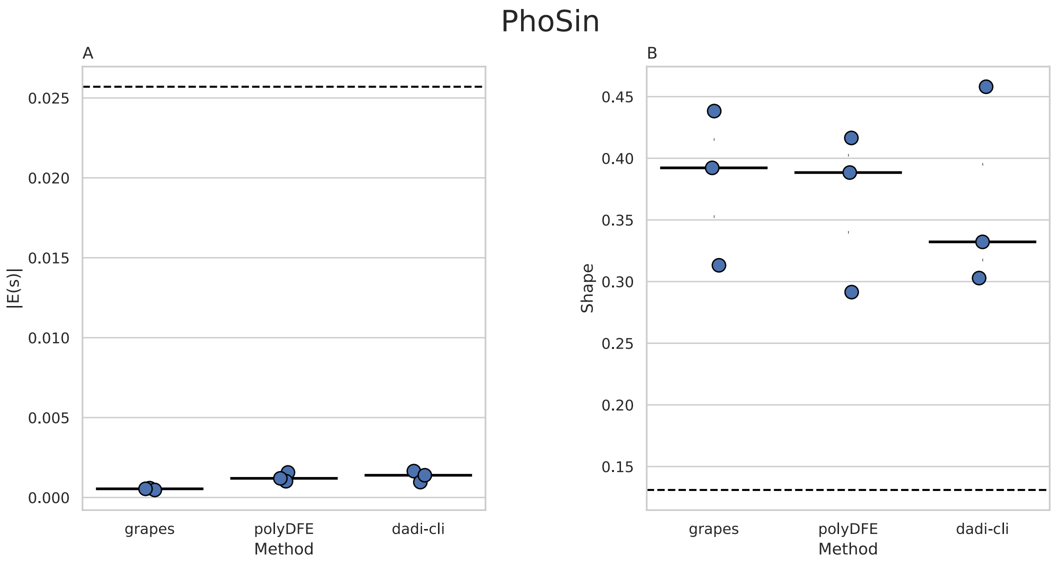


Figure S5: Inferred versus simulated DFEs from vaquita genomes simulated with a simple demographic model with one population with a constant size of $3,500$ and a gamma-distributed DFE acting on exons with a relationship between the selection coefficient ($s$) and dominance coefficient ($h$) (see **Methods**). DFE is inferred by three different methods: `GRAPES`, `polyDFE`, and `dadi-cli`. Mean absolute value of selection coefficient ($|E(s)|$, panel A) and shape parameter (panel B) are shown for each DFE inferred from all three simulation replicates with median values marked by horizontal bars and simulated values represented by dashed horizontal lines.

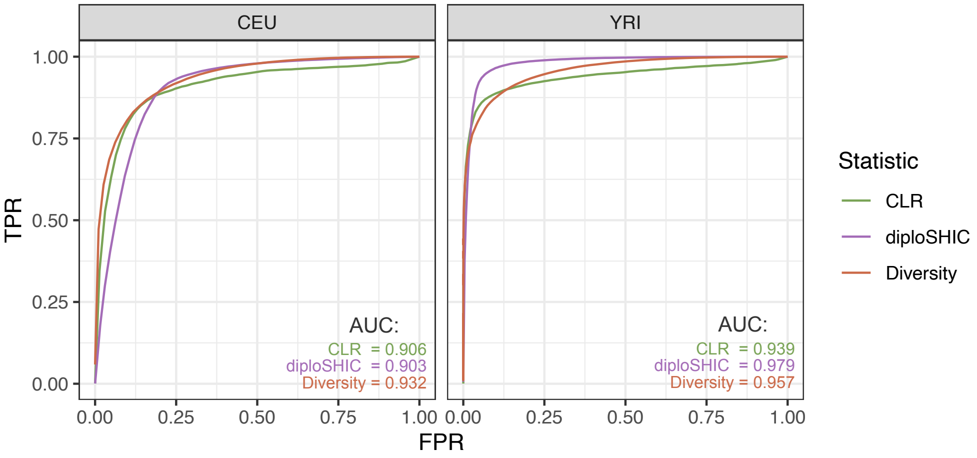


Figure S6: Receiver operating characteristic (ROC) curves for the three methods for detecting sweeps: `sweepfinder2` (labeled CLR), `diploshic`, and reduced diversity ($\pi$). True positive rates (TPR) were computed across all datasets simulated with selective sweeps, and false positive rates (FPR) were established by applying the methods to genomes simulated under the same settings but without sweeps (the BGS model; see **Methods**). Area under the curve (AUC) scores are shown for each method.

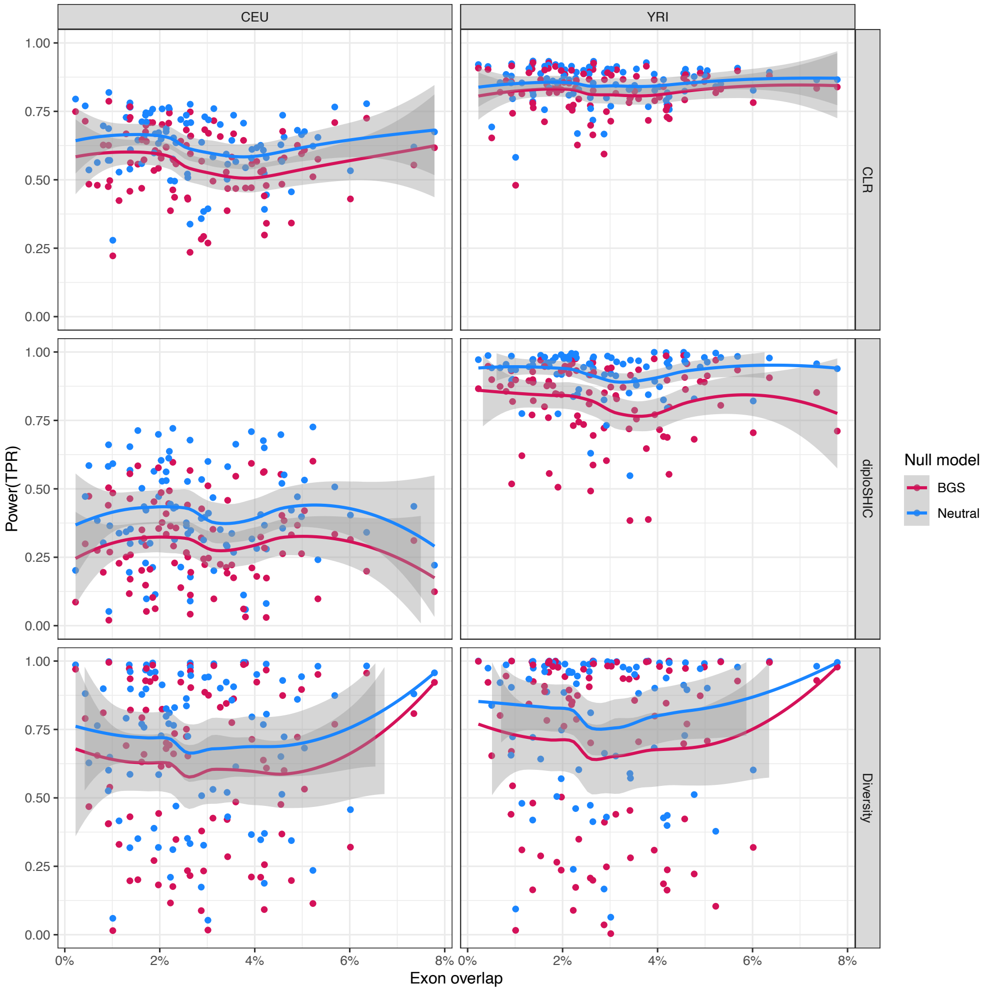


Figure S7: Power to detect selective sweeps as a function of local exon density. This figure shows the same power estimates shown in Figure 5, but with the genomic segments plotted against their exon density (percentage of exonic bases per window) instead of position along chromosome 1. Genomic segments were simulated with sweeps under a three population out-of-Africa model and with background selection from deleterious mutations in exons. Three methods for detecting sweeps were applied to simulated data: `sweepfinder2` (top row– labeled CLR), `diploshic` (middle row), and reduced diversity ($\pi$) (bottom row). Power (true positive rate) is shown for these methods for the CEU and YRI samples (left and right respectively). The thresholds of the test statistics were set to obtain a 5% false positive rate under a neutral null model (blue) and a null model with background selection from deleterious mutations in exons (red). Fitted lines represent loess smoothed regressions.

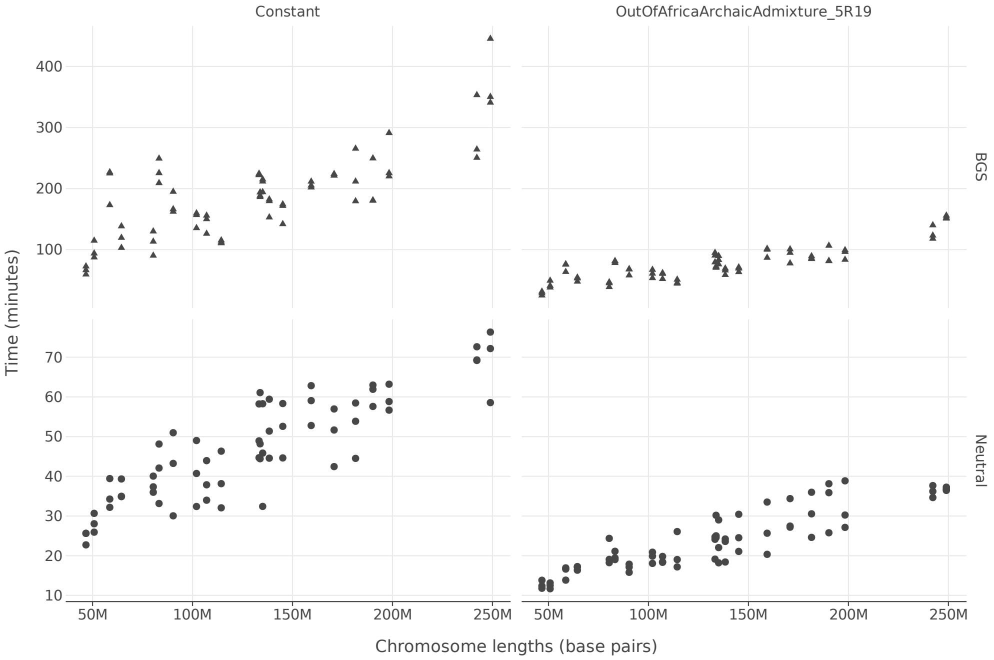


Figure S8: Simulation runtimes for human chromosomes with and without selection applied to exons and under two demographic models: a single population with constant size (left) and a three-population out-of-Africa model (right). Each point represents a single simulation of a single chromosome under conditions labeled by facets. Simulations were run on a mix of Intel and AMD machines (Intel Xeon Gold 6148 2.4 Ghz, Intel Xeon Gold 6248 2.5 Ghz, AMD EPYC 7513 2.6 Ghz and AMD EPYC 7713 2 Ghz).

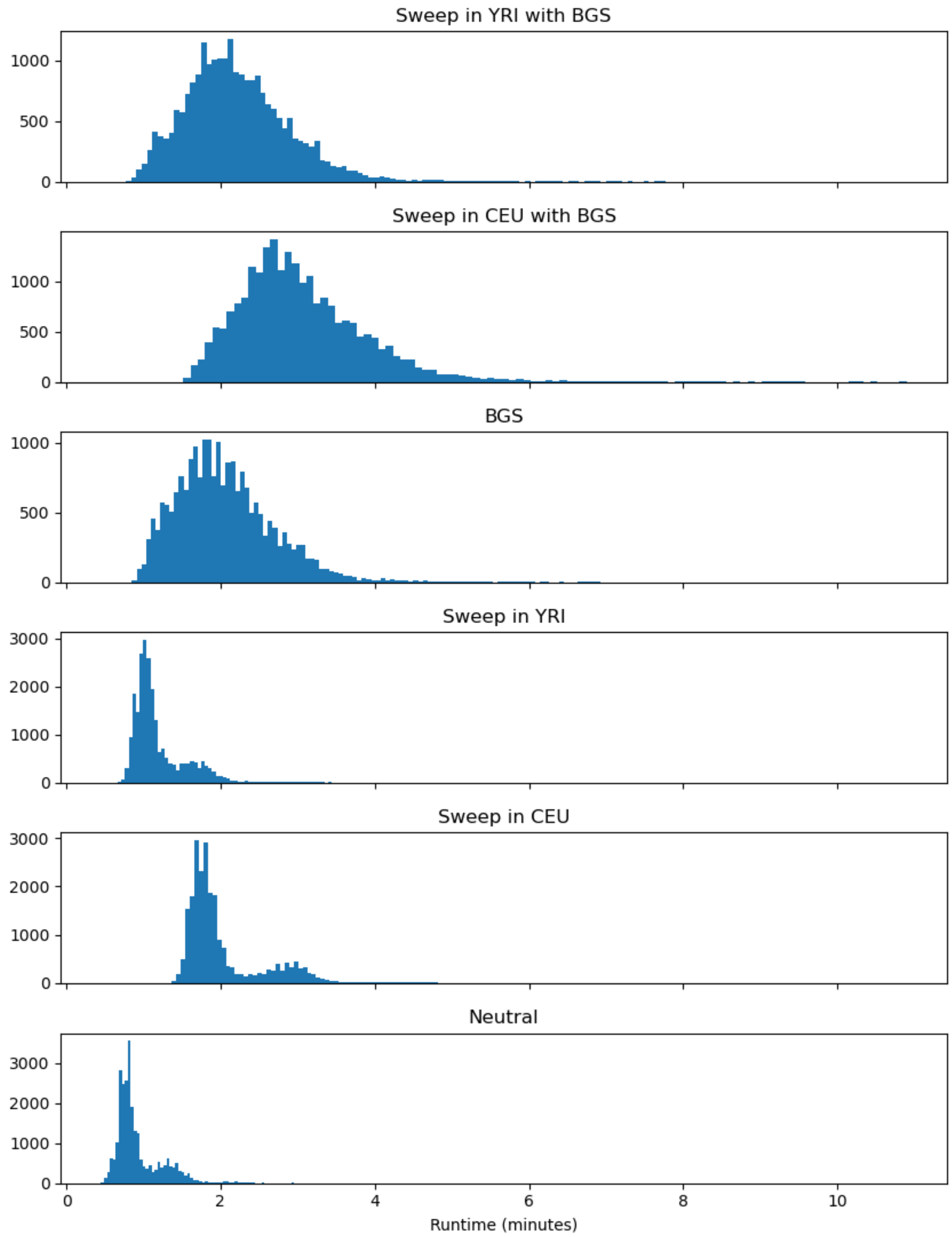


Figure S9: Distribution of simulation runtimes for segments of human chromosome 1 simulated for the sweep detection analysis. Runtimes are summarized separately for each of the six simulation conditions used in the analysis. Simulations were run on a mix of Intel and AMD machines (Intel Xeon Gold 6148 2.4 Ghz, Intel Xeon Gold 6248 2.5 Ghz, AMD EPYC 7513 2.6 Ghz and AMD EPYC 7713 2 Ghz).
